# Deciphering the Cofilin Oligomers via Intermolecular Disulfide Bond Formation: A Coarse-grained Molecular Dynamics Approach to Understanding Cofilin’s Regulation on Actin Filaments

**DOI:** 10.1101/2023.12.07.570543

**Authors:** Chengxuan Li, Tingyi Wei, Margaret S Cheung, Min-Yeh Tsai

## Abstract

Cofilin, a key actin-binding protein, orchestrates the dynamics of the actomyosin network through its actin-severing activity and by promoting the recycling of actin monomers. Recent experimental work suggests that cofilin also forms functionally distinct oligomers through thiol post-translational modification (PTM) that encourages actin nucleation and assembly. Despite these advances, the structural conformations of cofilin oligomers that modulate actin activity remain elusive because there are combinatorial ways to oxidize thiols in cysteines to form disulfide bonds rapidly. This study employs molecular dynamics simulations to investigate human cofilin 1 as a case study for exploring cofilin dimers via disulfide bond formation. Using the free energy profiling, our simulations unveil a range of probable cofilin dimer structures not represented in current Protein Data Bank entries. These candidate dimers are characterized by their distinct population distributions and relative free energies. Of particular note is a dimer featuring an interface between cysteines 139 and 147 residues, which demonstrates stable free energy characteristics and intriguingly symmetrical geometry. In contrast, the experimentally proposed dimer structure exhibits a less stable free energy profile. We also evaluate frustration quantification based on the energy landscape theory in the protein-protein interactions at the dimer interfaces. Notably, the 39-39 dimer configuration emerges as a promising candidate for forming cofilin tetramers, as substantiated by frustration analysis. Additionally, docking simulations with actin filaments further evaluate the stability of these cofilin dimer-actin complexes. Our findings thus offer a computational framework for understanding the role of thiol post-translational modification of cofilin proteins in regulating oligomerization, and the subsequent cofilin-mediated actin dynamics in the actomyosin network.

## INTRODUCTION

The actomyosin network serves as a critical cellular scaffold that regulates a variety of cell behaviors, including cell division, shape changes, and movement,^1,2^ regulated by actin-binding proteins. Among these numerous actin-binding proteins, cofilin holds particular significance. Cofilin monomers act as molecular scissors, severing actin filaments and facilitating the recycling process through the collaborative efforts of capping proteins, profilins, and other actin-binding proteins.^3–5^ When interacting with actin molecules, cofilin monomers employ distinct binding sites for globular actin monomers (G-actin) compared to actin filaments (F-actin), as highlighted in a previous study.^6^ A recent cryo-electron microscopy (cryoEM) investigation has elucidated the overall structure of F-actin with cofilin molecules decorating the filament’s surface, leading to the formation of what is termed cofilactin.^7^ These studies collectively suggest that filament surface heterogeneity plays a crucial role in modulating filament severing. The variation in filament severing boundaries arises from different combinations of cofilin-actin interfaces, contributing to the stochastic nature of filament severing.

Interestingly, experimental evidence substantiates that cofilin is not merely functional in its monomeric form; it also exists as oligomers with divergent roles in actin regulation.^8^ Cofilin has four cysteine (Cys) residues that can undergo redox-dependent post-translational modifications (PTM) to form specific intramolecular or intermolecular disulfide bonds.^9^ Specifically, rather than severing actin filaments, when cofilin forms oligomers through intermolecular disulfide bonds, they are implicated in promoting actin nucleation, assembly, and bundling.^10,11^ The formation and functioning of the cofilin oligomers are regulated by the local cofilin concentration and the phosphorylation pathway, which also governs the monomeric form.^11,12^ However, there are combinatorial ways to form intermolecular disulfide bonds. Establishing the causal relation between the molecular arrangement of cofilin oligomers and their distinct role in regulating actomyostin networks in response to redox is unclear.

Various experimental works have posited that these cofilin dimers likely emerge through intermolecular disulfide bonds between specific cysteine residues—namely, cysteine 39 and cysteine 147—on adjacent cofilin monomers,^8^ instead of a cofilin symmetric dimer through the disulfide bonds between their cysteine 139 residues.^13^ Moreover, prevailing theories suggest that the biologically active form of cofilin is not a dimer but rather a tetramer, leaving the dimeric state as a potentially transient state in oligomer formation.^11^ However, the structural and functional intricacies of these cofilin oligomers in response to thiol PTMs remain unknown, thereby offering ample opportunities for the application of coarse-grained simulation techniques.

While the atomistic simulations have shed light on crucial aspects of the mechanical stress from the composition of filament structures, they are often constrained by limited timescales and length scales, which pose challenges for achieving robust statistical analyses. Coarse-grained protein models provide an avenue to surmount these limitations. The application of coarse-grained protein modeling to explore the network dynamics of CaMKII/F-actin bundles, in conjunction with protein array experiments and electron microscopy imaging, has effectively elucidated the multivalent binding interactions between CaMKII and F-actin, aligning with experimental findings.^14^ Collectively, these studies underscore the capacity of coarse-grained protien models to capture sufficient molecular details, enabling them to replicate numerous features of filamentous behavior and, in turn, to address the regulatory roles of actin-binding proteins within the actin network.

The advantages of coarse-grained simulations in addressing complex protein behaviors over long timescales make them particularly well-suited for investigating the formation, stability, and function of cofilin oligomers and their interaction with actin filaments. In this study, we focus on human cofilin 1, with specific attention to the structural representation denoted by the PDB ID 4BEX.^13^ While it provides a symmetric cofilin dimeric structure comprised of two monomers, it is noteworthy that the authors themselves suggest that this assembly, formed during crystallization, may not faithfully represent a biologically relevant dimeric state. To fill the existing knowledge gaps, we provide a computational framework to evaluate the cofilin oligomers formed with distinct intermolecular disulfide bonds and their interaction with a short fragment of the actin filament.

## METHODS

### Associative-memory, water-mediated, structure and energy model (AWSEM)

The present study leveraged the AWSEM (Associative-Memory, Water-Mediated, Structure, and Energy Model) coarse-grained protein force field for an in-depth analysis of cofilin protein dimerization.^15^ This force field, implemented on the LAMMPS simulation platform, employs a three-bead model for each amino acid residue, representing C-alpha, C-beta, and O atoms. This coarse-graining preserves ideal peptide bond geometry, thus enabling efficient simulations of protein folding dynamics,^16^ structure prediction,^16^ and protein aggregation.^17–19^

The energy function of AWSEM comprises transferable and physically-motivated potentials, encapsulating complex residue-specific physicochemical properties such as hydrophobicity and electrostatic interactions within the framework of protein secondary and tertiary structures. Implicitly, the enthalpic contributions of protein-protein contacts are included in the AWSEM contact energies.

In AWSEM’s coarse-grained scheme, the energy function is sectioned into three main components: V_backbone_, V_non-backbone_, and V_FM_ (Fragment Memory), responsible for the backbone geometries, protein’s physicochemical attributes, and local structural tendencies, respectively. V_backbone_ encompasses five terms that regulate chain connectivity, bond angles around the C_α_ atom, orientations of the C_β_ atoms, backbone dihedral angles, and excluded volume interactions. Harmonic potentials control both the chain connectivity and the bond angles around the C_α_ atom. V_non-backbone_ is further split into three terms: V_contact_, V_burial_, and V_helical_. These terms individually consider aspects like tertiary fold contact interactions, residue exposure/burial preferences, and helical structure propensity. V_FM_ is tailored to bias the local structure towards those found in a “fragment memory” library of protein fragments with similar local sequences. This term also accounts for the local steric effects influenced by the protein’s local sequence. For this study, a single fragment memory scheme within AWSEM was employed to accentuate the effects of physical forces on folding and binding landscapes.

For further details on the force field, readers are referred to the work by Davtyan et al.^15^ The AWSEM code is publicly accessible via Github: AWSEM Repository (https://github.com/adavtyan/awsemmd).

### Structural preparation and simulation protocol for cofilin dimer formation through intermolecular disulfide bond

The wild-type sequence of human cofilin-1 (hCof1) is depicted in Figure 1A. Due to the limited availability of monomeric cofilin structures, a crystal structure with high resolution was selected for the monomeric form (PDB ID: 4BEX).^13^ This structure was found to possess a mutation at residue 147, where cystine was replaced by alanine. To revert the structure to its wild-type form, we mutated residue 147 back to cystine using the ‘Mutate Residue’ module available in VMD software, as shown in Figure 1B.^20^

**Figure 1.**
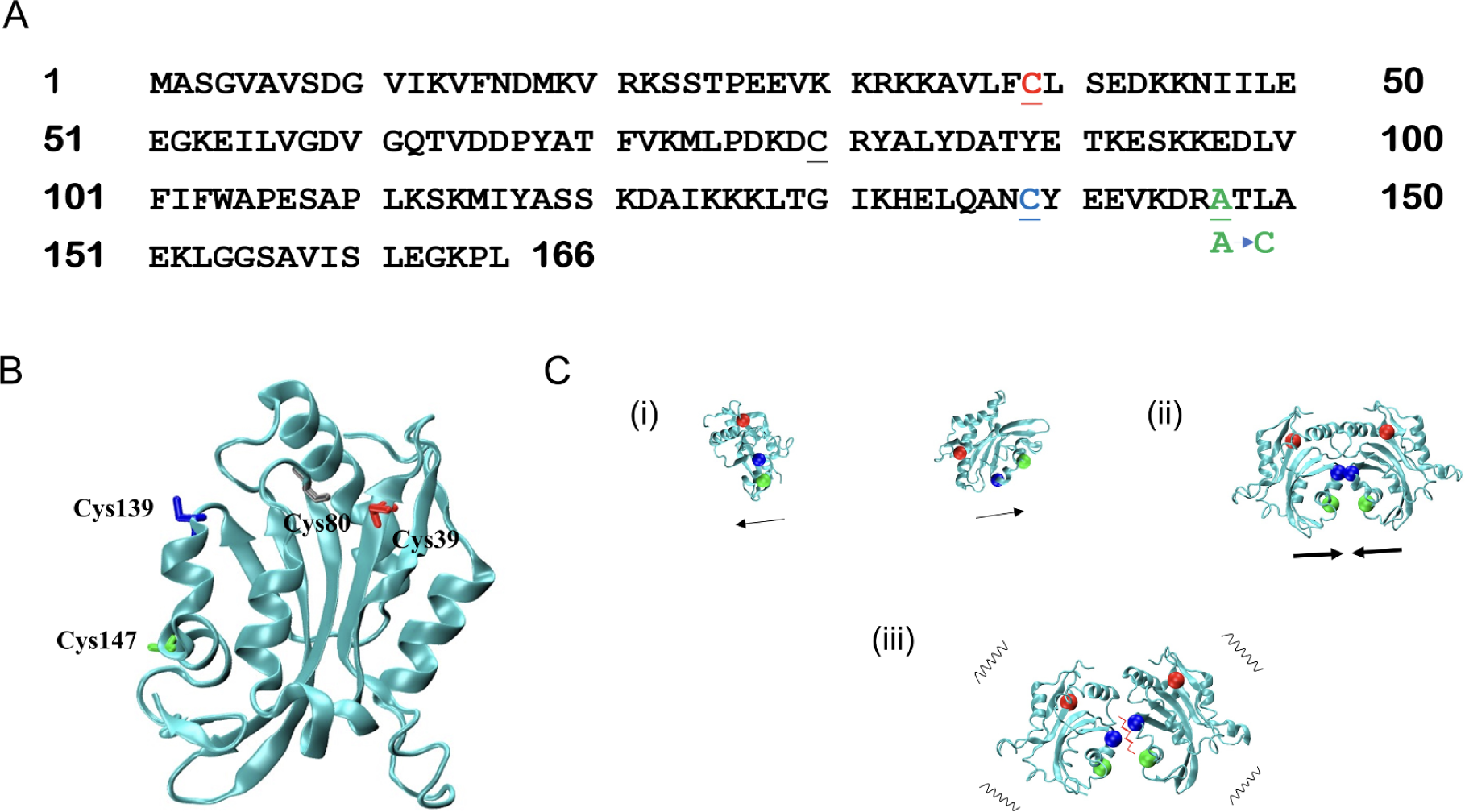
The mutated cofilin sequences, their monomeric structure, and simulated dimer formation workflow are described. A: Sequence representation of human cofilin 1 (PDB ID: 4BEX). Cysteine residues 39, 139, and 147 are highlighted in red, blue, and green, respectively. The latter was mutated to cysteine in our simulations to mirror *in vivo* conditions. While cysteine residue 80 is also underlined, it is not color-coded. The color patterns introduced here are consistently used in subsequent figures. B: Monomeric cofilin structure post the mutation at residue 147. The protein chain is rendered using the ’NewCartoon’ method in cyan, while cysteine residues employ the ’Licorice’ visualization with colors as designated in (A). Due to spatial proximity in relation to residue 39, cysteine residue 80 is labeled. C. The stages of simulated dimer formation. (i) Monomers are drawn apart using a gentle spring force centered on the total protein structures. (ii) The force from stage (i) is discontinued and replaced by a potent spring force — set at an equilibrium distance of 2Å, the approximate length of a disulfide bond — encouraging the monomers to assemble (as demonstrated using residues cys139 and cys139). (iii) Post removal of the bias from (ii), the monomers are at liberty to oscillate or dissociate. All molecular visualizations were made using VMD.

The simulation process for forming cofilin dimers was organized into three distinct steps, illustrated in Figure 1C: 1) Initiation (using pushing forces), 2) Association (using pulling forces), and 3) Relaxation (absence of biasing forces). In the Initiation step, a spring force characterized by an equilibrium distance greater than the intermolecular disulfide bond was employed to two cofilin monomers. During the Association phase, a spring force equivalent to the equilibrium distance of a disulfide bond was applied between the Cys39-Cys147 and Cys139-Cys139 residues of adjacent cofilin monomers, to simulate oxidized thiol cysteine residues from post-translational modifications. For simplicity, Figure 1C shows only the Cys139-Cys139 bond as an example. The Relaxation step was designed to allow the mutual configurations of cofilin monomers in the dimer assembly to stabilize naturally, simulating the reduced state of cysteine. Accordingly, the biasing spring force was removed during this phase.

The simulated system was held at a constant temperature of 300K throughout the study. Each simulation trajectory consisted of three phases: 1) Initiation, 2) Association, and 3) Relaxation, as previously outlined. These phases encompassed 0.5, 4, and 4 million simulation time steps, respectively. To ensure the robustness of our statistical analysis, we conducted a total of 30 individual trajectories.

### Importance sampling and free energy analysis

For the purpose of free energy analysis,^21^ we employed two distinct configurations of cofilin dimers as reference structures for umbrella sampling. The first configuration featured Cys139-Cys147 at the interface (139-147 cofilin dimer), which was identified during the previous unbiased relaxation simulation within the context of the 3-stage dimer formation simulation. The second configuration corresponded to the crystallized 4BEX assembly (139-139 cofilin dimer) obtained through experimental means^13^ Notably, the reference structure was derived by averaging over the last 50 simulation frames of a single trajectory, specifically chosen from among the 30 trajectories to ensure the stable dimer formation of interest.

In the umbrella sampling, a harmonic biasing force was applied to the α-carbon atoms of the target molecule, utilizing the global Q value of the entire dimer assembly as the sampling coordinate. The force constant was set at 1000 kcal/mol Å^2^, and the center values for each sampling window ranged from Q=0.025 to 0.975 at intervals of 0.025, yielding a total of 36 sampling windows. All other parameters were kept consistent with default values utilized in previous work.^16^

We conducted umbrella sampling analyses for five replicate sets, selecting only the binding simulations where the two cofilin monomers remained consistently associated with each other (i.e., Q ≥ 0.60). The global Q values from these binding simulations were concatenated into an array and subsequently categorized into 300 discretized centers. The data, now organized by centers, were processed through Pyemma thermo module to calculate the free energy and then visualized.^22^ The jupyter notebooks for umbrella sampling analysis can be found on the Github (https://github.com/pnnl/PTMPSI/tree/master/ptmpsi-awsem).

### Exclusion of long-range electrostatics in binding simulations

Recognizing the significance of electrostatic interactions in governing protein structure and dynamics is fundamental to our understanding of molecular behavior. Notably, while the standard AWSEM code incorporates local electrostatic interactions with the solvent, it does not consider long-range electrostatic interactions. Our previous research has emphasized the pivotal role played by long-range electrostatic interactions (namely, Debye-Hückel potentials) in predicting the structural characteristics of specific proteins and shaping the binding energy landscapes of various protein binders.^16^ However, when considering the stability of cofilin, these interactions appear to have a limited impact. A comparison of the simulated annealing results for cofilin, with and without the inclusion of long-range electrostatics, clearly demonstrates their negligible effect on the native structure’s stability, as visualized in Figure S1 in the Supporting Information.

Furthermore, even in scenarios where multiple cofilin molecules engage in dimerization, our frustration analysis (refer to Figure S2 in the Supporting Information) confirms that these long-range electrostatic interactions do not exert a significant influence. Figure S2 vividly illustrates the minimal role of electrostatic-induced frustration interactions at the cofilin binding interfaces. Consequently, despite the capability of AWSEM to incorporate long-range electrostatics in simulations, we have chosen to exclude these interactions from our binding simulations due to their limited impact on the proteins under investigation.

### Visualization, clustering, and frustration analysis

The analysis of our AWSEM simulations was conducted using various tools and techniques. To visualize the simulations, we employed VMD.^20^ The total protein structure was visualized using the ’New Cartoon’ drawing method, while the specific cysteine residues (39, 139, and 147) were highlighted using the ’Beads’ drawing method, each assigned a distinct color (Cys39 in red, Cys139 in blue, and Cys147 in green) for easy identification within the dimer.

For further trajectory analysis, we initially processed the PDB files containing the cofilin dimer configurations using the Python package MDTraj.^23^ Subsequently, we generated contact maps illustrating the residue positions within the cofilin dimers using the Contact Map Explorer (https://github.com/dwhswenson/contact_map), which is an open-source tool built on MDTraj. The scripts for generating these contact maps are available on our Github repository (https://github.com/pnnl/PTMPSI/tree/master/ptmpsi-awsem).

To perform structural clustering analysis, we curated a total of 1360 structures obtained from biased simulations conducted for umbrella sampling. In pursuit of a comprehensive exploration of the cofilin dimer ensemble, we systematically selected these structures across five separate replicate simulations, each representing 16 distinct trajectories (resulting in a total of 5x16=80 trajectories). Within each trajectory, we extracted a structure every 50 simulation frames, yielding a grand total of 5x16x17=1360 representative structures. Our clustering analysis began by transforming the Q-global data, into a square form. The Q value serves as a structural similarity measure, ranging from 0 to 1, where “1” signifies an identical reference structure and “0” denotes a completely dissimilar counterpart. Subsequently, we employed hierarchical clustering analysis, utilizing the Python package Seaborn and the function “seaborn.clustermap.” In this analysis, we specified critical parameters, including metric=“correlation” for the pairwise distance metric and method=“average” for the linkage method. These choices facilitated the generation of informative visualizations. Please consult our GitHub repository (https://github.com/pnnl/PTMPSI/tree/master/ptmpsi-awsem) for further reference and access to detailed scripts related to the clustering analysis of the dimer configurations.

To investigate frustration within the cofilin structures, we utilized the Frustratometer Server, accessible at http://frustratometer.qb.fcen.uba.ar/.^24^ The concept of frustration in biomolecules and its contemporary perspective on folding, function, and assembly have been discussed in detail in other sources.^25,26^ This analysis was performed with a sequence separation parameter of 3 and without considering electrostatics (see **Exclusion of electrostatics in binding simulations** section above).

## RESULTS

### AWSEM simulations reveal multiple probable binding interfaces in a cofilin dimer

In our AWSEM simulations, we identified multiple cofilin dimer candidates established by intermolecular disulfide bond with distinct binding interfaces, of which six were particularly noteworthy. Cofilin’s mutual configurations in the dimer are represented using three key cysteine residues (39, 139, 147) for visual inspection. Previous experimental findings had proposed a dimer interface involving cysteine 39 and 147 residues, referred to as the 39-147 cofilin dimer.^10^ Additionally, the PDB structure of human cofilin 1 (PDB ID: 4BEX) reported a disulfide bond formed between adjacent, symmetric monomers at cysteine 139 residues, known as the 139-139 cofilin dimer.^13^ While our simulations confirmed the presence of the 39-147 cofilin dimer, the experimentally reported 4BEX assembly involving disulfide bonds between cysteine 139 residues was notably absent. Our simulations also unveiled several previously unreported binding interfaces, with the 39-39 and 139-147 interfaces being the most prevalent. In Figure 2A, the population distribution of these cofilin dimer candidates is illustrated using structural clustering analysis, while Figure 2B visualizes the centroid structures based on their constituent cysteine residues. It is important to note that the population size shown in Figure 2A serves as a qualitative indicator for structural categorization purposes and does not represent the realistic population of representative clusters.

**Figure 2.**
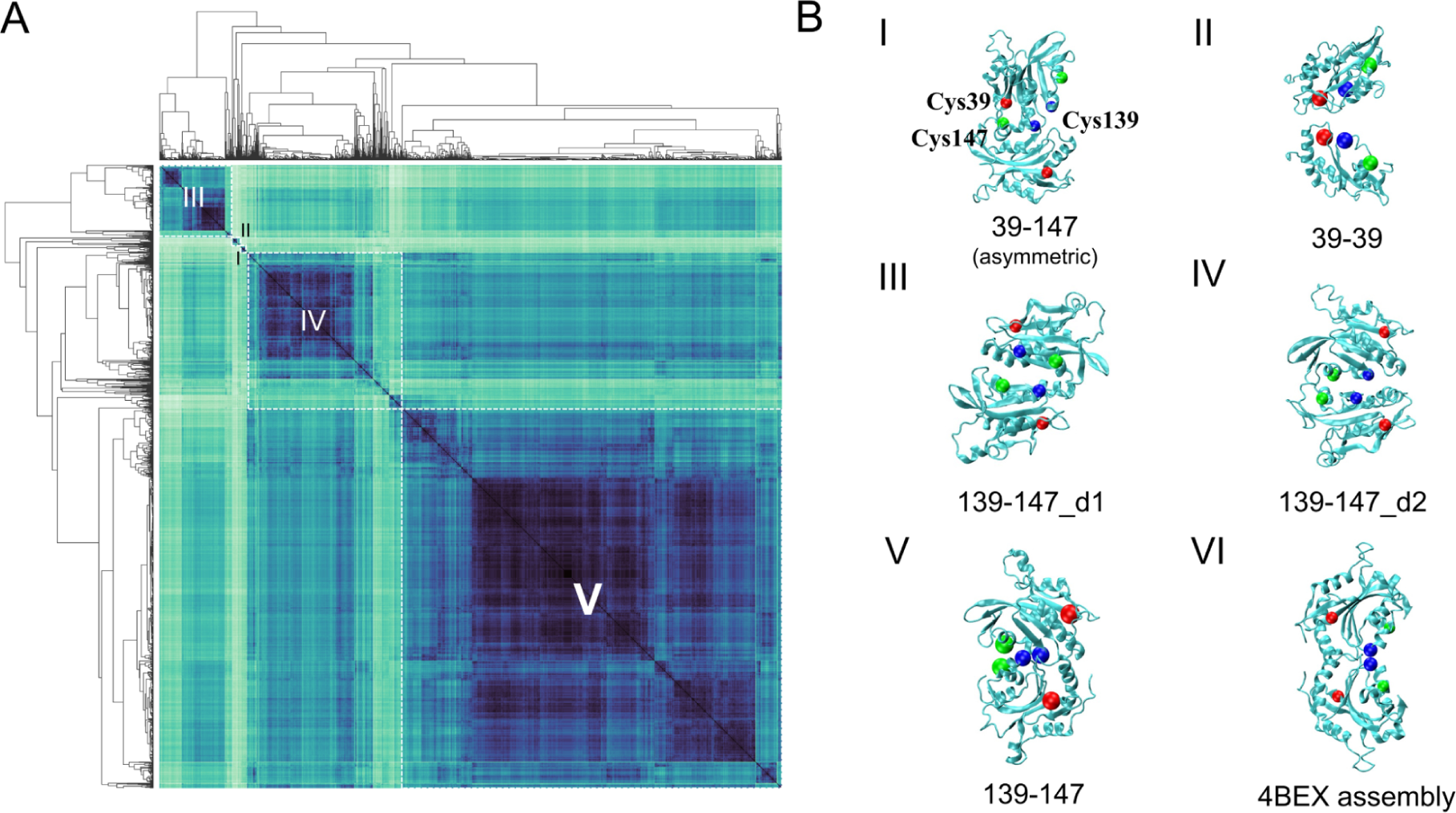
Analysis of Structural Clustering for Various Cofilin Dimer Configurations is shown. A. Clustered heatmap with dendrograms of cofilin dimeric configurations derived from AWSEM MD simulation trajectories using the Q value for structure. Dark clusters correspond to the cofilin dimer candidates depicted in B. Cofilin dimer candidates indexed I-V, correlating with the cluster map in A. Candidate VI (139-139) represents the experimental assembly from 4BEX, which was not identified in our simulations. Candidates I-V are denoted by the cysteine residues detected at the interface. The cysteine residues 39, 139, and 147 are represented as red, blue, and green spheres, respectively. Candidates III and IV (both belonging to 139-147) are further distinguished as d1 and d2, representing degenerative states 1 and 2, respectively. Notably, the candidate I (39-147) dimer exhibits asymmetry, while the other candidates are symmetric.

Our simulation results predict that the 39-147 cofilin dimer is the only asymmetric assembly among the other six configurations. The asymmetry of this assembly can be visualized in three-dimensional space by representing the three key cysteine residues (39, 139, and 147) with red, blue, and green balls, respectively (refer to Figure B - I). One can conceptualize a virtual triangle formed by connecting these three cysteine residues (colored balls) within the cofilin molecule. The asymmetric binding configuration of a cofilin dimer is akin to envisioning two virtual triangles stacked together with their vertices misaligned. This asymmetric assembly suggests a generic intermolecular interaction between cofilin molecules *in vivo* and is known to exhibit bundling activity under oxidative stress.^10^ The interface residue pairs in this asymmetric assembly consist of a combination of mild hydrophobic contacts (involving Ser-Ile, Ser-Cys, Tyr-Cys, Cys-Gly, Leu-Gly) and scattered charge pairs (including Glu-Lys, Glu-Glu).

The experimentally determined cofilin assembly, the 139-139 cofilin dimer, exhibits C2 symmetry, with two virtual triangles mirroring each other in the crossing plane. In this symmetric assembly, the interface residue pairs engage in numerous strong hydrophobic contacts (e.g., Phe-Leu, Ala-Leu, Met-Met, Ala-Met, Ala-Ala, Cys-Cys, Cys-Tyr, Asn-Pro) and a significant number of repulsive charged pairs (e.g., Glu-Glu, Lys-Lys). Despite its high degree of symmetry, the distribution of the resulting interaction pairs does not optimize at the interface, thus remaining unobserved in our simulations.

On the other hand, the predicted 139-147 cofilin dimer displays considerable conformational dynamics among its subspecies. These subspecies share a significant number of strong hydrophobic contacts (e.g., Met-Leu, Ala-Leu, Leu-Ile, Met-Met, Met-Ala, Cys-Cys) along with mild stabilizing charge-pairs (e.g., Asp-Glu). This combination of chemical interactions underscores the conformational flexibility and dynamic exchange of contacts at the interface.

Interestingly, the model of 39-39 cofilin dimer exhibits a highly symmetric molecular structure (C2 symmetry) with very limited hydrophobic contacts, nearly none at all. The majority of the binding interface is composed of charged residues such as Lys(+), Glu(-), and Asp(-), resulting in charge-complementary clusters, including major attractive charge-pairs, Lys(+)-Glu(-) and Lys(+)-Asp(-), and several repulsive charge-pairs like Glu(-)-Glu(-) and Glu(-)-Asp(-). This phenomenon highlights a unique charge-pair distribution that compensates for repulsive charge forces through effective distribution. One thing worth mentioning is that Arg(+) is not observed as part of the electrostatics-driven binding interface, which is notable considering its common role in electrostatic interactions. One possible explanation for the exclusion of Arg(+) at the binding interface is the interplay between steric considerations, the structural specificity required for protein-protein interactions, and the relatively large and complex side chain of Arginine (Arg) compared to some other charged residues like Lysine (Lys). Protein-protein interactions necessitate a precise fit between interacting surfaces, and the choice of residues in the binding interface often aims to achieve optimal interactions, prevent steric clashes, and maintain a specific structural arrangement.

### Free Energy Landscapes of Major Cofilin Dimer Candidates: Insights from Importance Sampling

Despite the plethora of cofilin dimer configurations obtained through clustering analysis, their thermodynamic stability remains unknown. To assess their relative stability among the six cofilin dimers, we employed the umbrella sampling technique to explore the binding free energy landscape of cofilin dimers. Figure 3 presents the free energy profile of cofilin dimer candidates using the 139-147 dimer as the reference structure. Notably, several basins are evident along the free energy profile, indicating stable populations of the dimer. These free energy basins are labeled A&B, C, and D, with their respective representative structures (dimer candidates I to VI) displayed accordingly.

**Figure 3.**
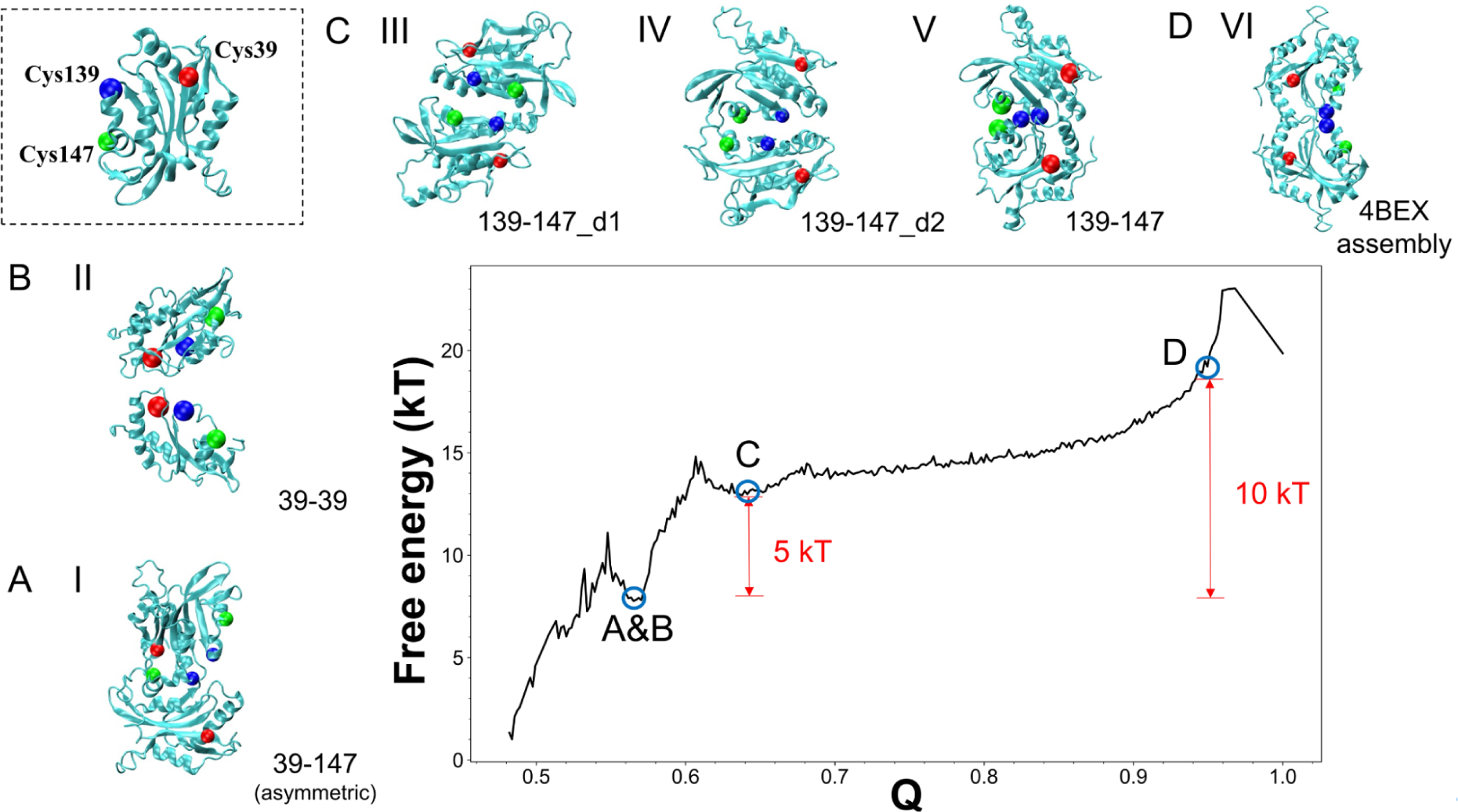
Free Energy Profiles of Cofilin Dimer Candidates are shown. The monomeric cofilin structure is illustrated at the top left: the main protein chain is in cyan, while cysteine residues 39, 139, and 147 are colored red, blue, and green, respectively. This color scheme extends to all dimer structures within this figure. A, B, C, and D denote various free energy profiles, either basin-like or non-basin-like, corresponding to specific dimer configurations. A. Candidate I: 39-147 dimer. B. Candidate II: 39-39 dimer. C. Candidates III-V: 139-147 dimers along with their degenerative states. D. Candidate VI: 139-139 dimer (4BEX experimental configuration). Notably, Candidates I and II exhibit a local free energy minimum that is 5 *kT* less than Candidates III-V and 10 *kT* lower than Candidate VI. Candidate VI does not present a local free energy minimum.

Dimer candidates I (39-147) and II (39-39) are situated within the A&B basin, representing the lowest free energy state among the others, despite their initial low prevalence as observed in the clustered dendrogram. This result suggests a greater thermodynamic stability of approximately 5kT (∼3kcal/mole at 300K) compared to our other dimer candidates, namely III, IV, and V (e.g., the 139-147 dimer). The 139-147 dimer ensemble is characterized by the free energy basin C, which exhibits a broader range, indicating higher conformational dynamics. Consequently, the 139-147 dimer encompasses several degenerate configurations, including 139-147 itself and a doublet pair, d1/d2. One distinctive feature that distinguishes d1 from d2 is the relative placement of Cys139 and Cys147 within the interface. In the case of d1, the positioning of one cofilin’s Cys139 and Cys147 in the interface is on opposite sides compared to those of the other cofilin. Conversely, d2 demonstrates that Cys139 and Cys147 are situated on the same side. This side-by-side variation is primarily attributed to a rotational change at the interface. Alongside this rotational difference, it’s worth noting that they also exhibit similar global Q values, as illustrated in the two-dimensional free energy landscape depicted below. The 139-139 dimer (VI) is not stable in our simulation as it does not exhibit any free energy basin, suggesting it is not a biologically relevant dimer (in line with the statement in the paper^13^).

### Experimentally characterized cofilin assembly exhibits a highly frustrated interface while predicted dimer models exhibit a less frustrated interface

We conduct frustration analysis^24^ for the aforementioned set of six cofilin dimer candidates (see Figure 4). Frustration in the context of protein analysis refers to energetically unfavorable interactions within specific regions of a given protein. Frustration analysis typically involves comparing the energy of a given interaction or region with the statistical energy distribution of different decoy states.^25,27,28^ If the energy of the interaction is higher than the average energy of the decoys, it is considered frustrated (indicated by red lines). This signifies that the local interaction is not stable within the current protein structure. Conversely, if the energy of the interaction is lower than that of the decoys, it is termed minimally frustrated (indicated by green lines), implying that the local interaction energy cannot be further stabilized through mutation or state changes. Please refer to the Methods section for a detailed definition of frustration and a quantitative description. The dimer interfaces of candidates I, II, and V exhibit predominantly green lines with few instances of red lines, indicative of minor frustrated interactions as compared to candidates III and IV. This observation suggests that the interfaces of candidates I, II, and V are favorable binding counterparts. However, candidate VI, corresponding to the 4BEX experimentally determined assembly, displays many red lines in its frustration analysis, implying a high degree of frustrated interactions. Consequently, the candidate VI interface is unlikely to be a favorable binding interface in the biological context.

**Figure 4.**
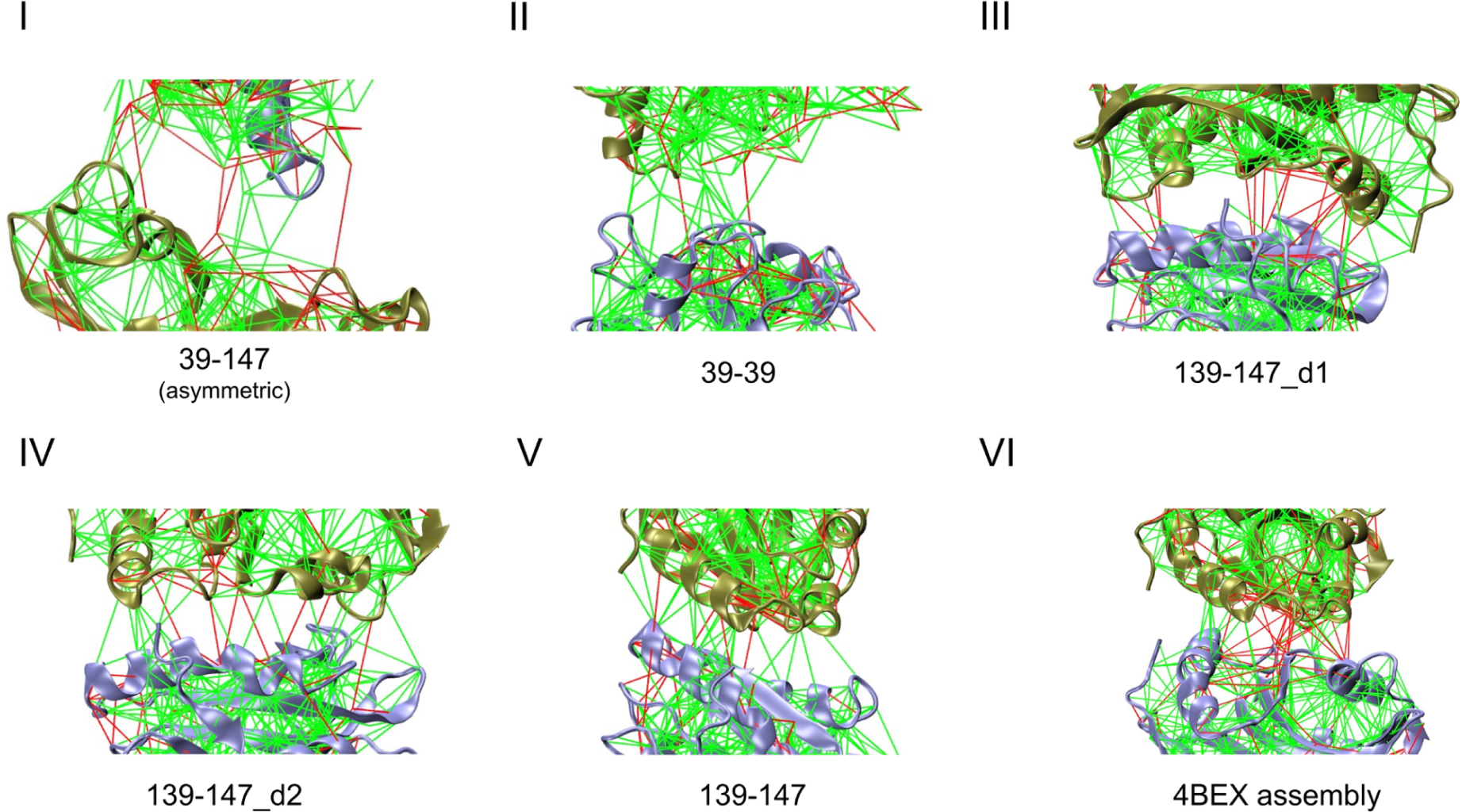
Frustration Analysis of Cofilin Dimer Interfaces is shown. Interactions at the binding interface across various cofilin dimer candidates are analyzed for frustration. Green lines highlight minimally frustrated residue pairs, while red lines highlight highly frustrated residue pairs. Notably, Candidates I, II, and V demonstrate less frustrating interfaces compared to Candidates III and IV. In contrast, Candidate VI displays pronounced frustration at the binding interface.

### The population shifts of dimeric structures from I to V are dynamic due to low free energy barriers in between basins

In the context of relatively low free energy barriers separating the basins populated with predicted cofilin dimer configurations, we employed importance (umbrella) sampling simulations to construct two-dimensional free energy landscapes. These landscapes profile free energy as functions of critical coordinates. Figure 5 illustrates these landscapes, depicting the free energy in terms of the global Q value and the radius of gyration (Figure 5A), as well as the system’s potential energy as a function of the radius of gyration (Figure 5B). Thermodynamically speaking, candidate V readily interconverts with candidates III and IV, owing to negligible energy barriers. It’s noteworthy that both III and IV exhibit comparable Q values, approximately around Q=0.6, within the same free energy basin. This similarity in Q values indicates that they share similar contacts formed at the interface. This similarity in the interface suggests a degree of dynamic flexibility inherent to this binding interface. It is characterized by variations in the relative orientations of individual monomers, which results in a degeneracy in energy, designated as d1 and d2. In contrast, dimer configuration V presents a significantly distinct Q value, approximately 0.9, indicating a structurally different configuration. Remarkably, the radius of gyration (R_g_) for III-V falls within the range of 20 to 22 Å, consistent with their similarity in shape. Furthermore, these local minima are more thermodynamically favorable for conformational transitions into candidates I and II, as indicated by the directional arrows in Figure 5.

**Figure 5.**
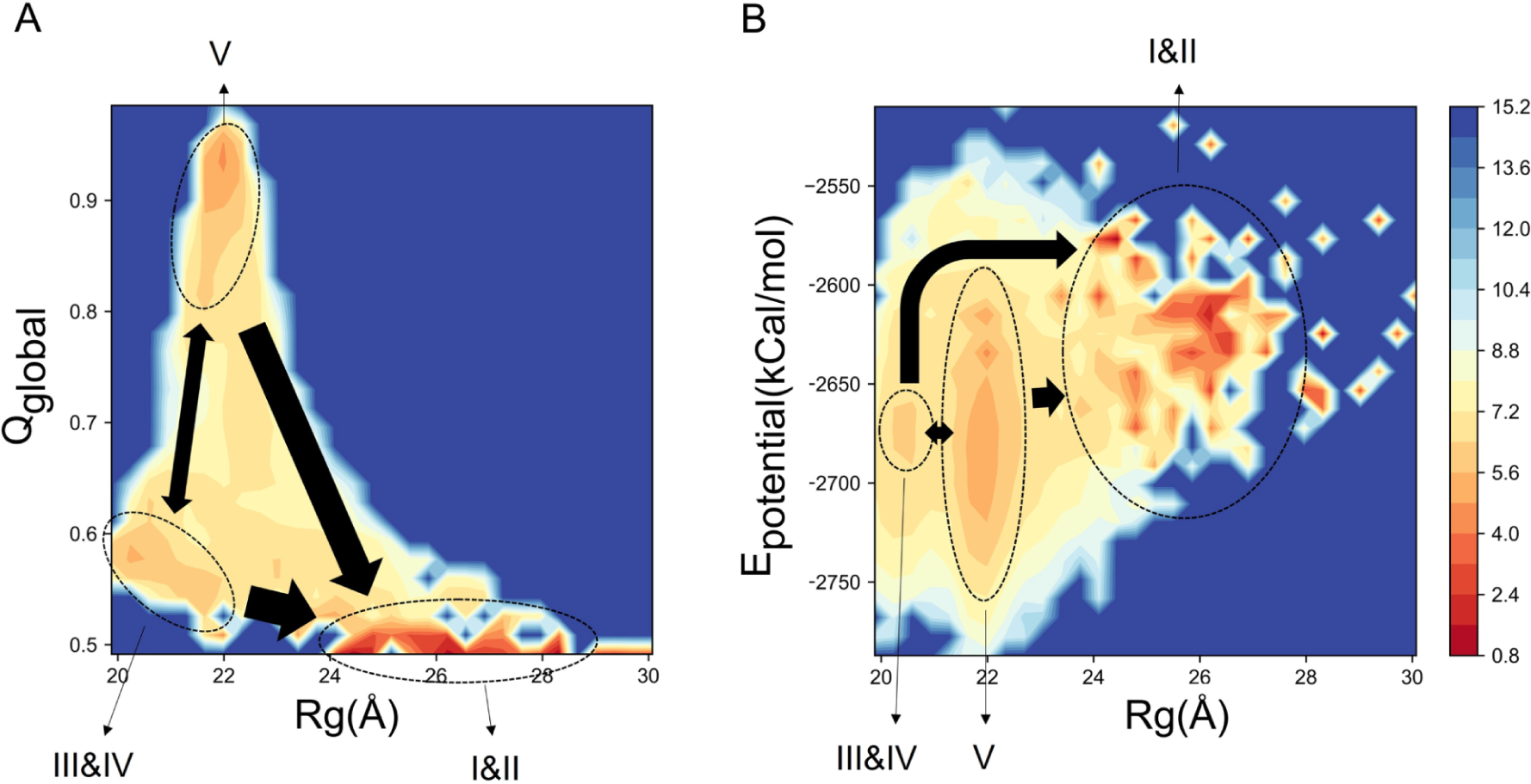
Free Energy Landscape of Cofilin Dimer Transitions is explored using two-dimensional free energy surfaces. (A) Depiction of the two-dimensional free energy surface based on the radius of gyration (R_g_) and the global Q value (with reference to candidate V). (B) Free energy landscape mapped as a function of R_g_ and potential energy. Dimers are pinpointed at the free energy local minima, represented by dashed circles, and labeled I to V. Arrows of varying thickness indicate the probability of transitions, influenced by relative free energy differences.

### Cofilin monomer and dimers interact differently with actin filament fragments

To explore the binding affinities of various cofilin forms with actin, we utilized the ClusPro2.0 server for docking simulations between a cofilin dimer model and a fragment of F-actin. As a control, we first demonstrated that monomeric cofilin readily docks onto a F-actin fragment where the cofilin monomer (yellow) places at the junction of actin units, as depicted in Figure 6A—an observation consistent with its role in actin severing. Conversely, cofilin dimers (for example Model I) can only dock at the terminal ends of F-actin, not between adjacent actin units, as illustrated in Figure 6C. Figures 6B and 6D further illustrate that both monomeric and dimeric forms of cofilin exhibit strong binding affinity for G-actin. Based on these findings, we propose that the smaller size of the cofilin monomer facilitates its docking between actin units on F-actin, thus enhancing its role in severing. Conversely, the larger size of dimeric cofilin limits its binding to either G-actin or the terminal ends of F-actin, aligning with their respective roles in actin nucleation and assembly.

**Figure 6.**
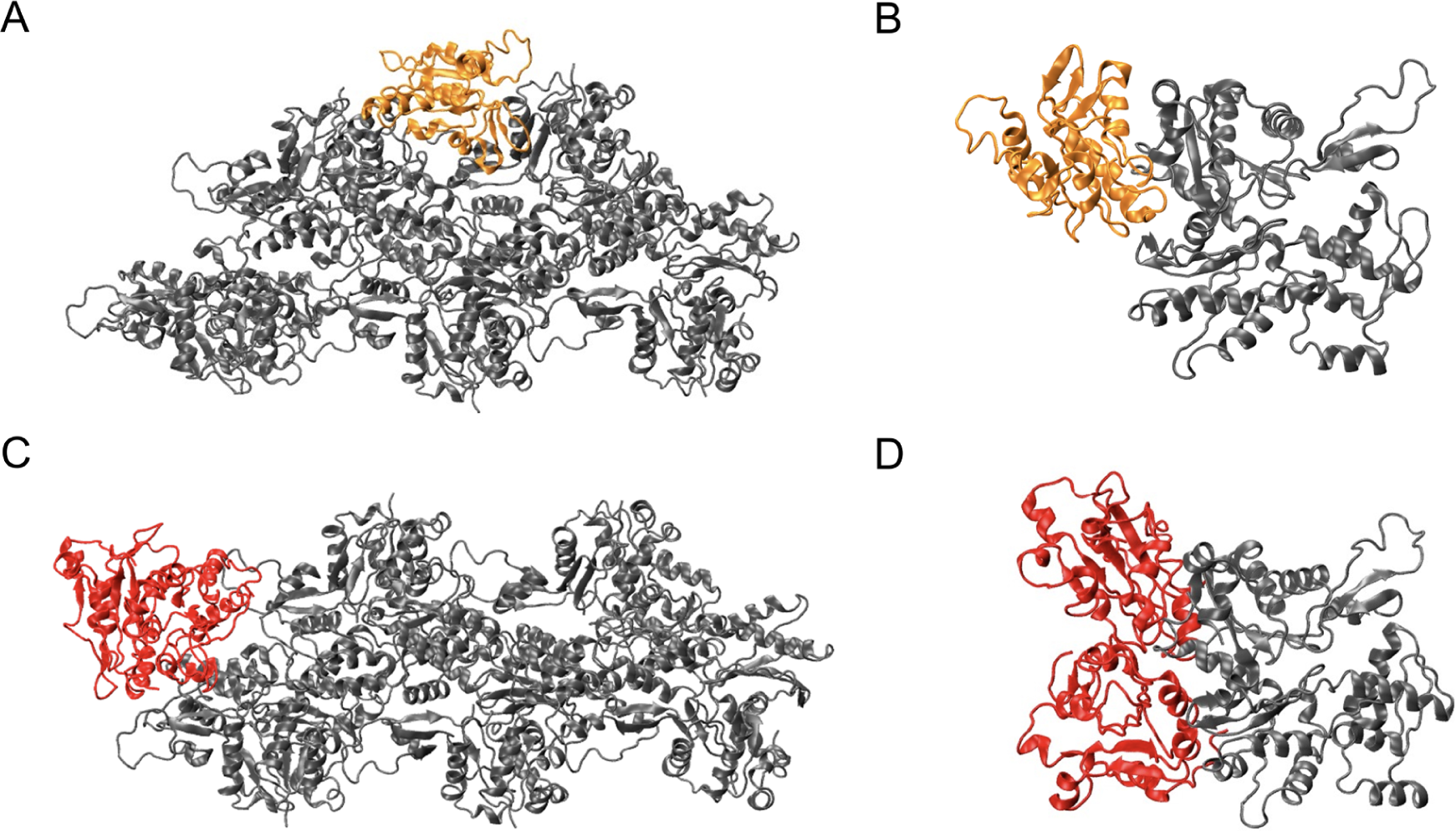
Interaction Dynamics of Cofilin with Actin Structures is predicted and shown. (A) Interaction of a monomeric cofilin (displayed in orange) with a filament-actin (F-actin) structure (shown in gray), which consists of 5 actin subunits. (B) Monomeric cofilin (orange) interacts with a G-actin structure (gray). (C) Dimeric cofilin structure (colored in red) docked onto an F-actin structure (gray), composed of 5 actin units. (D) Interaction of a dimeric cofilin (red) with a G-actin (gray). Visualization of the docking process of cofilin monomers and dimers onto filament-actin is facilitated by the ClusPro server.

## DISCUSSIONS

### Symmetry and Stability of Cofilin Dimers: Insights to Cofilin Tetramers from AWSEM Simulations

In this simulation study, we have predicted various cofilin dimer configurations that could potentially be guided by intermolecular disulfide bond formation. We should note that each cofilin dimer is characterized by distinct binding interfaces. Employing free energy analysis^29^ through the AWSEM coarse-grained force field,^15^ these proposed dimeric configurations display diverse population distributions and relative free energies. Among these, one particular candidate, designated as “dimer 139-147,” stands out due to its prevalence, characterized by an interface involving cysteine residues at positions 139 and 147. Additionally, we identify two other notable dimer candidates named “dimer 39-39” and “dimer 39-147.” The configuration of “dimer 39-147” bears a resemblance to the experimentally proposed dimer structure, while “dimer 39-39” holds promise as a potential precursor to cofilin tetramer formation.

Some of these configurations find support in experimental evidence, either directly or indirectly, while others are theoretical predictions lacking experimental validation. Nevertheless, our free energy calculations have demonstrated the capacity to predict the stability of these configurations. Notably, among all the predicted cofilin dimer configurations, the 39-39 assembly stands out due to its remarkable stability, high degree of symmetry, and optimized charge-pair distributions at the interface. This prediction raises the intriguing possibility of a prevalent population when cofilin oligomerizes into larger entities -- from cofilin dimers to tetramers, where two cofilin dimers randomly diffuse and collide.

The 39-39 dimer’s C4-like symmetry positions it as a strong candidate for participating in the formation of a cofilin tetramer. To test this hypothesis, we conducted AWSEM simulations. Initially, we applied two biasing forces to maintain disulfide bond equilibrium distances between the Cys39 residues of adjacent 39-39 dimers. Subsequently, these forces were removed to assess the tetramer’s structural stability. As illustrated in Figure 7A, these simulations revealed a stable cofilin tetramer configuration. Further analysis of mutational frustration within the tetramer, as shown in Figure 7B, indicated minimal frustration at the interfaces between the four chains. We speculate that this tetrameric configuration is energetically favorable, thereby providing support for its potential as a biologically relevant assembly. While higher-order reaction schemes are possible, we consider the most probable reaction channel to involve a second-order reaction, wherein two activated cofilin dimers interact to form a tetramer. Such simulations will require more sophisticated simulations; therefore, it is beyond the scope of this study.

**Figure 7.**
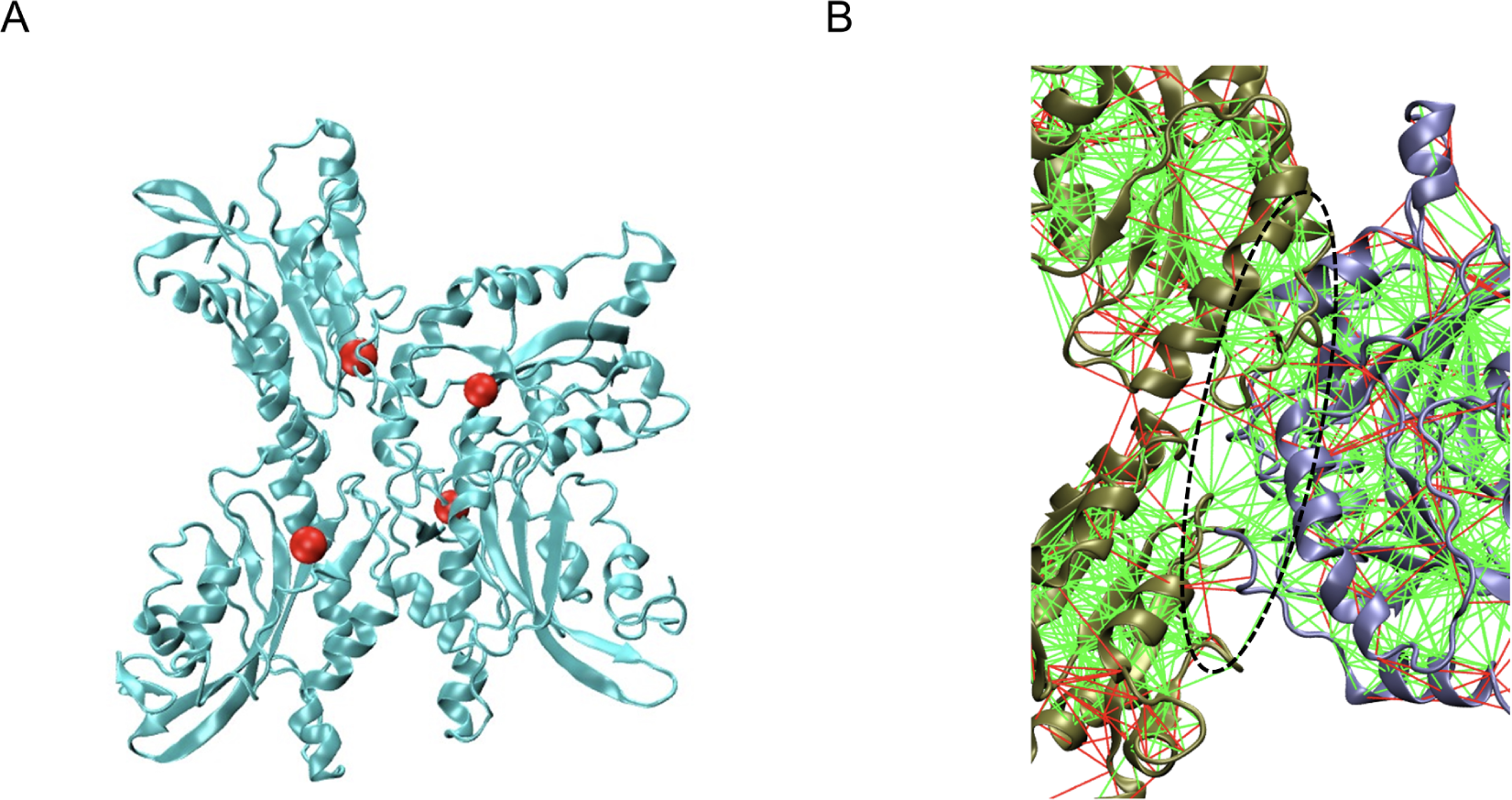
The Proposed Tetrameric Configuration of Cofilin Using 39-39 Dimers is shown. (A) Display of the simulated tetrameric cofilin structure, with protein chains rendered in cyan and cysteine 39 residues emphasized using red balls. Notably, this configuration remains stable in the AWSEM simulation without any external force biases. (B) Mutational frustration analysis was conducted at the interface between two contrasting dimers, colored tan and gray. A dominant minimally frustrated interface is inferred, as the majority of interactions within the delineated ellipsoid are green, indicative of an optimized binding interface.

### Cofilin Oligomerization and Its Functional Implications in Actin Regulation through Post-translational Modifications

The regulatory influence of cofilin monomers or oligomers on filamentous actin, for instance, involves a complex interplay of cooperative binding and mechanical coupling with the filament, resulting in diverse severing activities at the interfaces between bare and decorated segments.^30^ Numerous computational investigations, conducted at various levels, have been dedicated to exploring the mechanical stress modulation induced by cofilin binding.^31–33^ Among these investigations, mesoscopic models have proven to be particularly valuable for elucidating the mechano-elastic properties of actin filaments in the context of fragmentation. De La Cruz et al. pioneered a mesoscopic model capable of predicting bending and torsional rigidities based on polymer interaction energies, geometric constraints, and twisting-bending coupling dynamics.^31^ This model, as subsequently refined to include considerations of filament helicity and occupancy, has demonstrated its efficacy in mapping filament strain energy across specific lateral and longitudinal interfaces.^33^ Since these mechanical stress requires input from subtle changes in protein structures, all-atom molecular dynamics simulations play a pivotal role in furnishing these intricate structural insights at the molecular level that remain elusive in mesoscopic models. For instance, extensive molecular dynamics simulations have the capability to faithfully reproduce the subtle helical twists observed in actin filaments upon cofilin binding,^34^ a phenomenon consistent with cryo-electron microscopy findings.^35^ The atomistic-scale simulations have also uncovered the asymmetric dynamics of barbed and pointed ends, along with intricate conformational changes in neighboring subunits arising from differences in crossover lengths at slow-severing and fast-severing boundaries.^36,37^

Even with AlphaFold2^38^ or RoseTTAFold^39^ that offers us a tool to predict protein structures from sequences accurately, these structure-only models with canonical amino acids lack the interpretation of protein functions that rely on post-translational modification (PTM). PTM is the chemical modification of amino acids in response to changes in the cellular environment. While some PTMs, such as phosphorylation, require specific enzymes to characterize cellular behaviors, oxidation-reduction (redox)-based PTMs (i.e., redox PTMs) of thiol-containing cysteines are the most common in rapid responses to shifting redox conditions.^40^ One of the authors recently developed a Python-based workflow, *PTM-Ps*i (A Python Package to Facilitate the Computational Investigation of <u>P</u>ost-<u>T</u>ranslational <u>M</u>odification on <u>P</u>rotein <u>S</u>tructures and Their <u>I</u>mpacts on Dynamics and Functions), that allows users to interpret how chemical perturbations caused by PTMs, particularly thiol PTMs, affect a protein’s properties, dynamics, and interactions with its binding partners. They demonstrated the utility of *PTM-Psi* for interpreting sequence-structure-function relationships of a cysteine-rich protein derived from thiol redox proteomics data.^41^

Here, cofilin oligomerization involves thiol PTMs of cysteines to form intermolecular disulfide bonds under oxidation stress. Our cofilin dimer models enable us to explore these possibilities based on cofilin’s distinct binding interfaces. We employed a straightforward docking approach to address these questions, aiming to provide practical insights into predicting cofilin’s function in terms of its oligomeric state. We showed that due to the volume exclusion, only cofilin monomers could fit at the cleft of actin filaments, while cofilin dimers have fewer options to bind to the actin filament. Under oxidation stress, cofilins may regulate their binding with actin filament through its thiol PTMs of disulfide formation. We fit this coarse-grained modeling module based on AWSEM simulations into the schema of *PTM-Psi* (https://github.com/pnnl/PTMPSI/tree/master/ptmpsi-awsem).

## CONCLUSIONS

Our study focuses on cofilin, a vital actin-binding protein known for its role in actin-severing and monomer recycling. Recent experimental discoveries have highlighted cofilin’s capacity to form functionally diverse oligomers through intermolecular disulfide bond formation, contributing to actin nucleation and assembly. The formation of cofilin oligomers signifies its importance in regulating actin dynamics under oxidative stress. We provided a computer model to evaluate the structural conformations of these cofilin oligomers and evaluated their impact on actin binding with coarse-grained molecular simulations. Our findings enhance our comprehension of the intricate interplay between cofilin and actin, which has a far-reaching impact on understanding the cellular cytoskeletal dynamics.

## ACKNOWLEDGEMENT

This work was supported by the National Science and Technology Council (NSTC), Taiwan, Grant No. 110-2113-M-194-009-MY2. MSC is grateful for the support from the National Science Foundation PHY 2019745 and MCB 2221824. We appreciate our interactions over many years with the Wolynes group at Rice University on several implementation details of AWSEM and especially the developments made by Weihua Zheng while at Rice.

