## Supplementary information for "Deciphering the Cofilin Oligomers via Intermolecular Disulfide Bond Formation: A Coarse-grained Molecular Dynamics Approach to Understanding Cofilin’s Regulation on Actin Filaments"

**Supporting Information** 1

**Deciphering the Cofilin Oligomers via Intermolecular Disulfide Bond Formation: A Coarse-grained Molecular Dynamics Approach to Understanding Cofilin's Regulation on Actin Filaments** 1

Figure S1 - Effects of long-range electrostatic interactions on the structural stability of human cofilin 1 are evaluated using AWSEM. 2
Figure S2 - Single residues frustration analysis 3
Figure S3 - 4BEX-assembly mutational frustration analysis 5
Figure S4 – 39-39 mutational frustration analysis 7
Figure S5 - 39-147 mutational frustration analysis 9
Figure S6 - 139-147 mutational frustration analysis 11
Figure S7 - Comparison of the four cofilin dimers' mutational frustration analysis is shown. 12

**(a)**

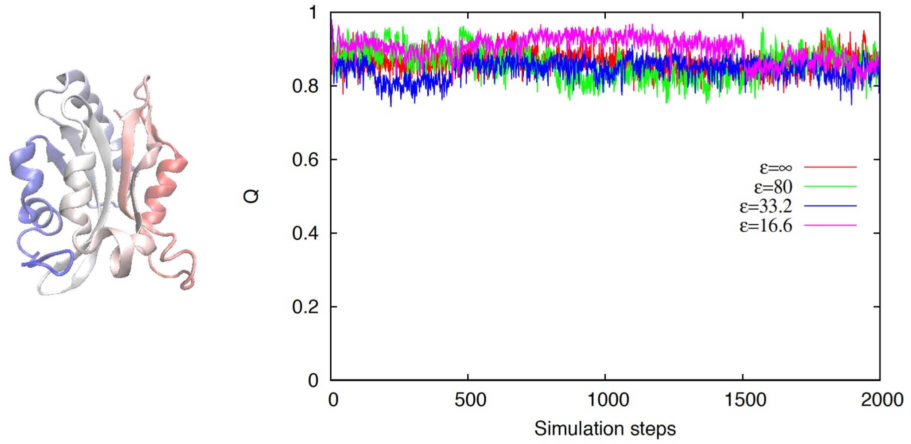


**(b)**

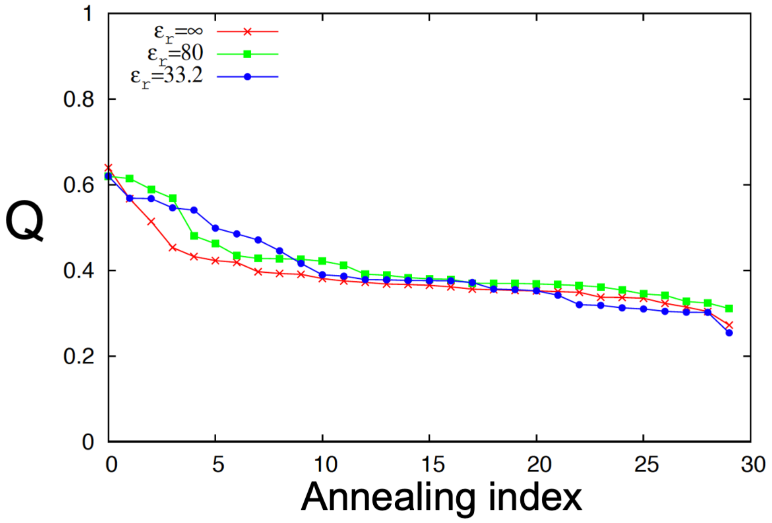


Figure S1 - Effects of long-range electrostatic interactions on the structural stability of human cofilin 1 are evaluated using AWSEM.

(a) Structural stability test. The Q value of cofilin is calculated as a function of simulation time. The Q value remains 0.8~0.9 (high Q value), independent of the strength of long-range electrostatics. (b) Simulated annealing profiles. The annealing result shows similar profile, irrespective of the electrostatic strengths. The Debye-Huckel potentials were used to mimic the long-range electrostatic effects. ε represents the dielectric constant with ε=∞, 80 (water), 33.2, and 16.6 meaning "no", "mild", "strong", and "extreme strong" electrostatic strength, respectively.

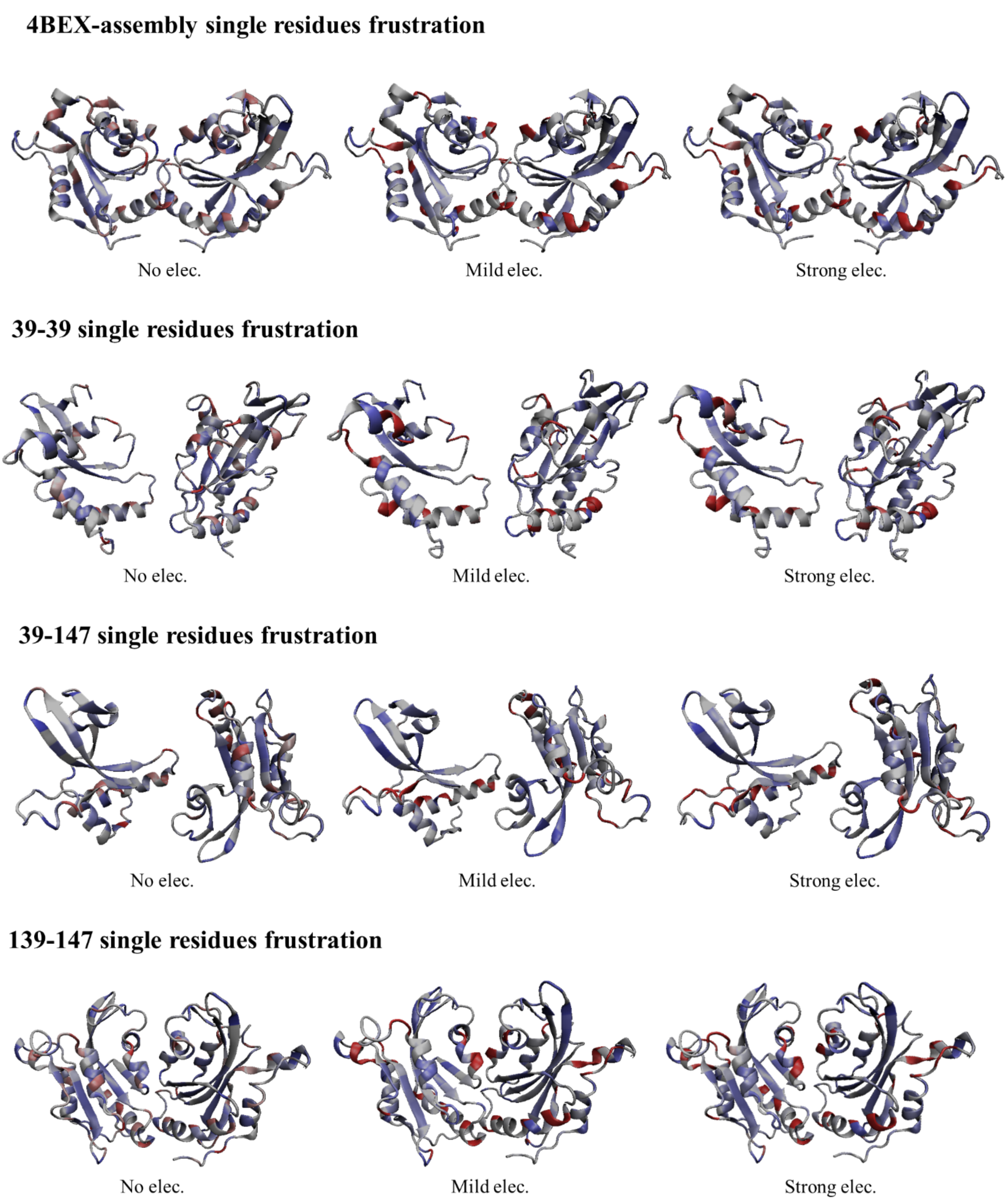


### Figure S2 - Single residues frustration analysis

Electrostatically-induced frustration is evaluated using single residue frustration mode for four cofilin dimer configurations (From top to bottem: 4BEX experimental assembly, 39-39 dimer, 39-147 dimer, 139-147 dimer). The frustration is calucated (from left to right) in the context of "No electrostatics (No elec.)", "Mild electrostatics (Mild elec.)", and "strong electrostatics (Srong elec.)", which represents dielectric constant in the Debye Hückel potentials $\varepsilon = \infty$, 80, 33, respectively. The cofilin structure is shown using cartoon diagram with highly frustrated residues colored in red, neural residues colored in gray, and minimally frustrated residues colored in purple.

No elec.

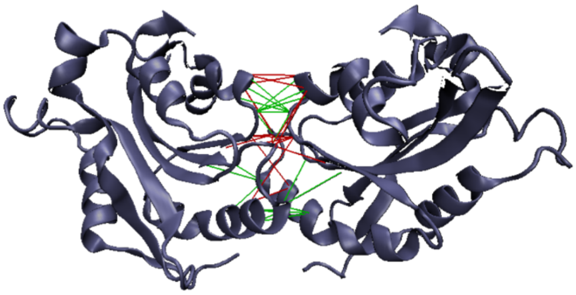
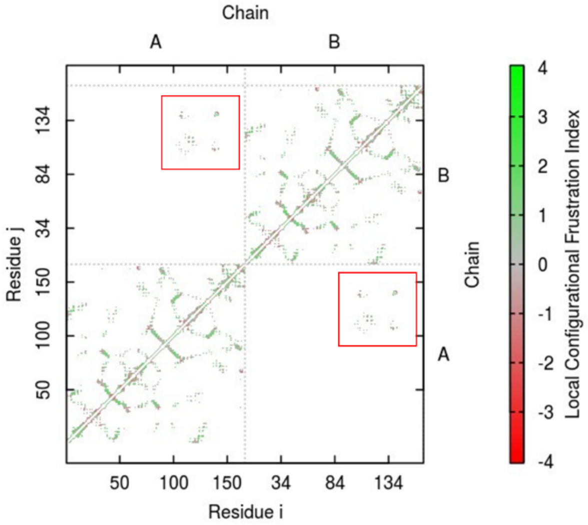


Mild elec.

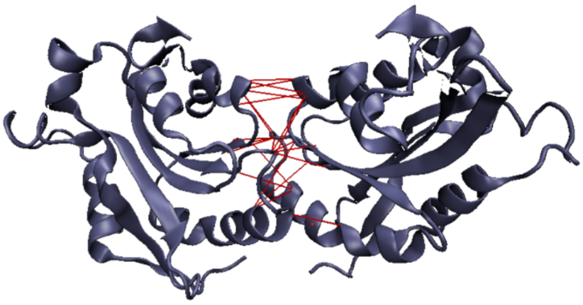
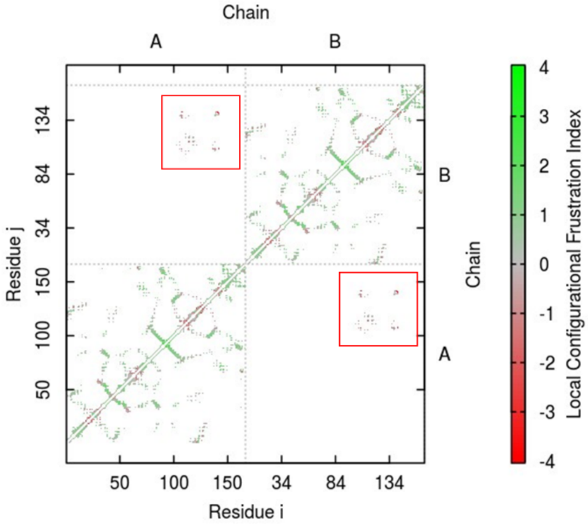


Strong elec.

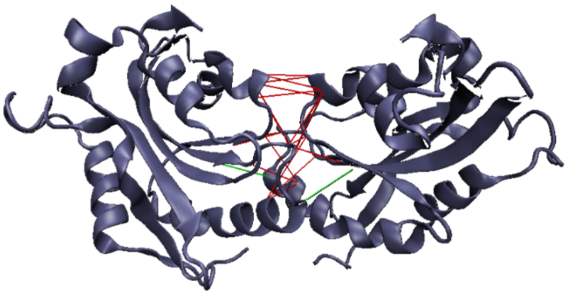
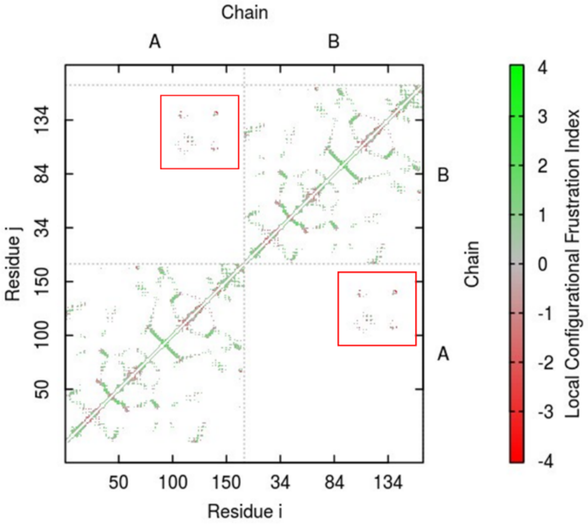

### Figure S3 - 4BEX-assembly mutational frustration analysis

Visualization of the 4BEX dimer (left) and the corresponding contact map (right). In the contact map on the right, red boxes represent interface residues in the dimer on the left. Frustration calculations are performed with varying electrostatic conditions labeled as "No elec.", "Mild elec.", and "Strong elec." (from top to bottom), corresponding to dielectric constants $\varepsilon = \infty$, 80, and 33, respectively (in Debye-Hückel potentials). The cofilin structure is depicted in a cartoon diagram, with highly frustrated residues shown in red and minimally frustrated residues in green. As electrostatic strength increases, interface interactions become more frustrated (indicated by the red line).

No elec.

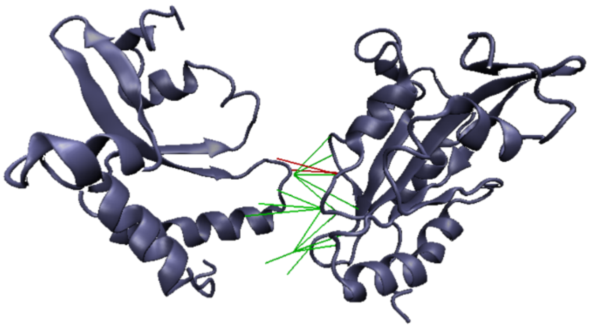

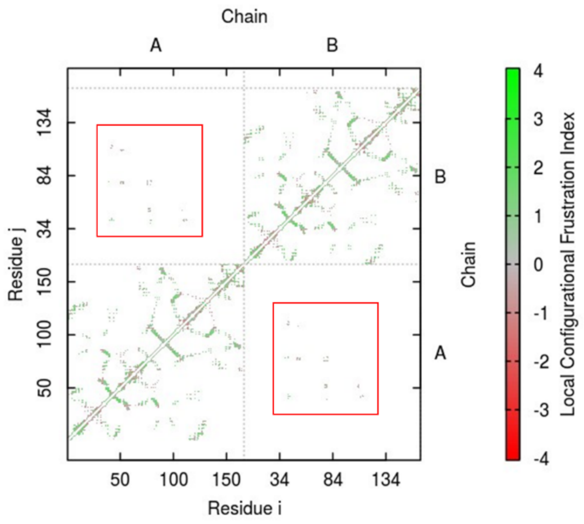


Mild elec.

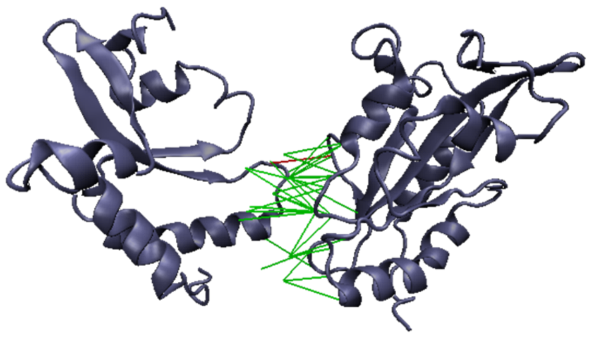

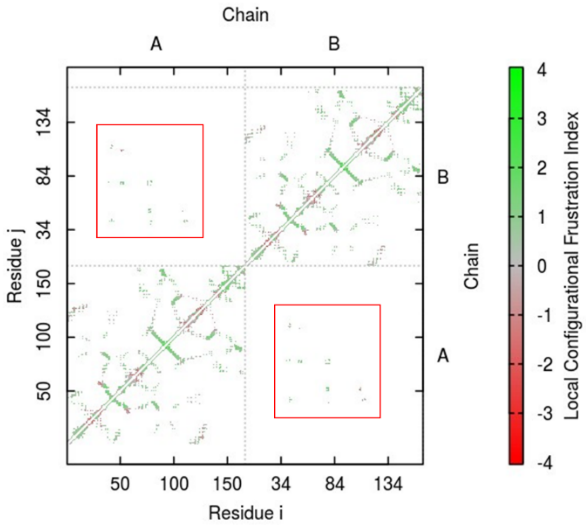


Strong elec.

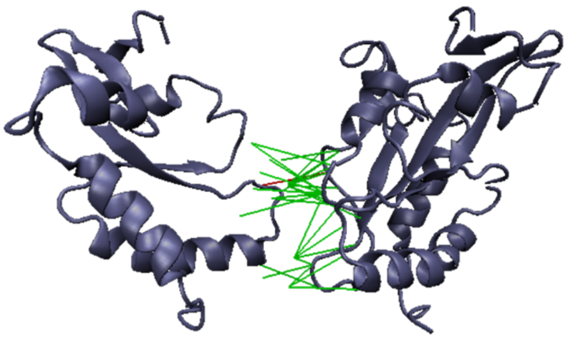

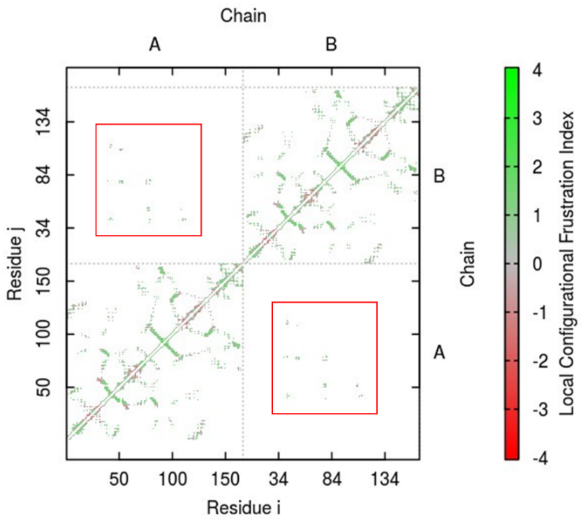

### Figure S4 – 39-39 mutational frustration analysis

Visualization of the 39-39 dimer (left) and the corresponding contact map (right). Red boxes in the contact map on the right indicate interface residues in the dimer on the left. Frustration analysis conducted under varying electrostatic conditions labeled as "No elec.", "Mild elec.", and "Strong elec." (from top to bottom), with dielectric constants $\varepsilon = \infty$, 80, and 33, respectively (in Debye-Hückel potentials). The cofilin structure is represented in a cartoon diagram, with highly frustrated residues highlighted in red and minimally frustrated residues in green. In comparison to the 4BEX-assembly dimer, the 39-39 dimer displayed minor changes in the proportion of frustrated (red line) and minimally frustrated (green line) residues under different electrostatic forces.

No elec.

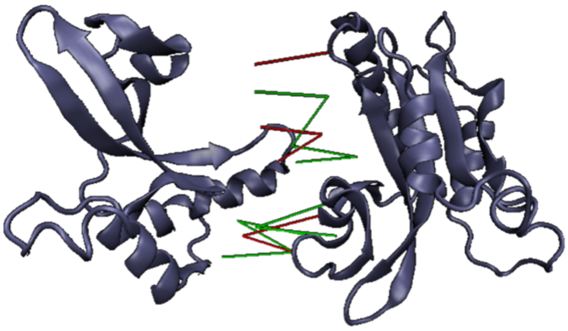

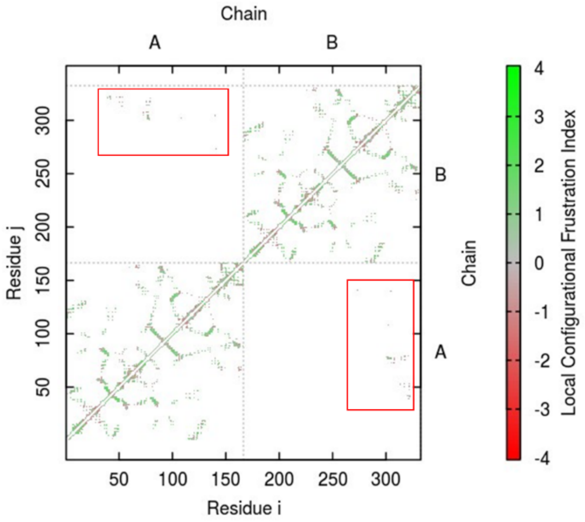


Mild elec.

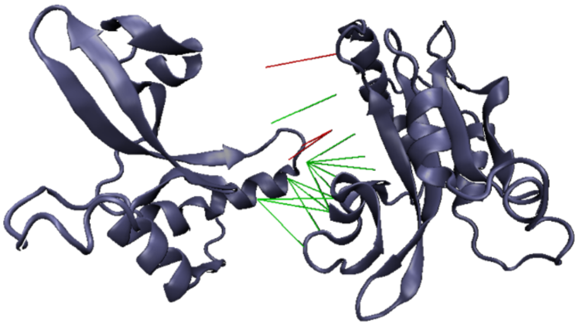

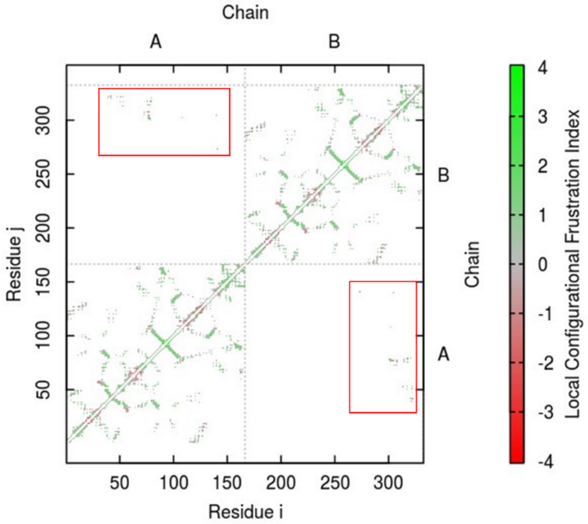


Strong elec.

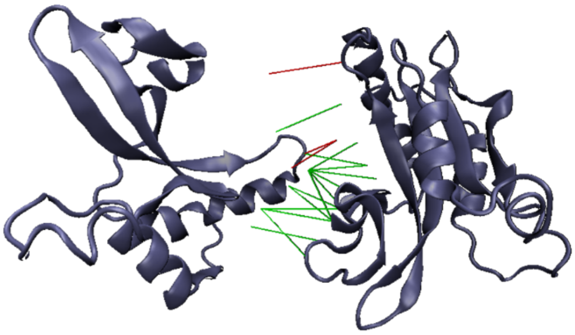

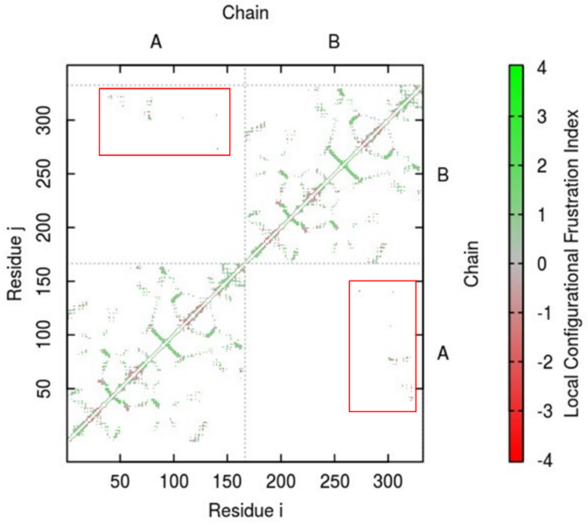

### Figure S5 - 39-147 mutational frustration analysis

Visualization of the 39-147 dimer (left) and the corresponding contact map (right). Red boxes in the contact map on the right indicate interface residues in the dimer on the left. Frustration analysis conducted under varying electrostatic conditions labeled as "No elec.", "Mild elec.", and "Strong elec." (from top to bottom), with dielectric constants $\varepsilon = \infty$, 80, and 33, respectively (in Debye-Hückel potentials). The cofilin structure is represented in a cartoon diagram, with highly frustrated residues highlighted in red and minimally frustrated residues in green. The 39-147 dimer exhibits consistent proportions of frustrated and minimally frustrated residues under the influence of the three electrostatic forces, similar to the 39-39 dimer.

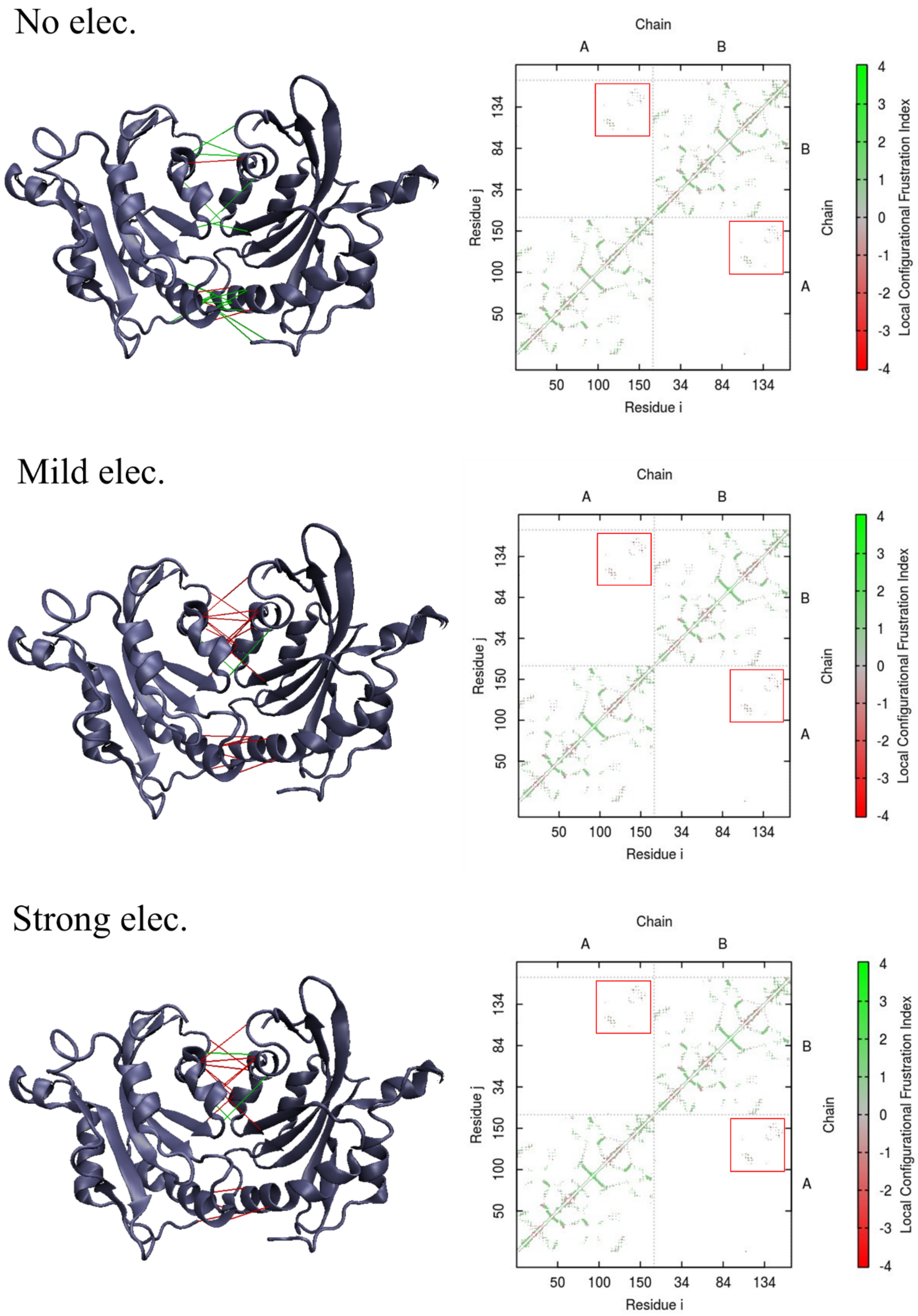
No elec.
Mild elec.
Strong elec.
Chain
A
B
Residue j
Residue i
50
100
150
34
84
134
Local Configurational Frustration Index
4
3
2
1
0
-1
-2
-3
-4

### Figure S6 - 139-147 mutational frustration analysis

Visualization of the 139-147 dimer (left) and the corresponding contact map (right). Red boxes in the contact map on the right highlight interface residues in the dimer on the left. Frustration analysis performed under different electrostatic conditions denoted as "No elec.", "Mild elec.", and "Strong elec." (from top to bottom), corresponding to dielectric constants $\varepsilon = \infty$, 80, and 33, respectively (in Debye-Hückel potentials). The cofilin structure is depicted in a cartoon diagram, with highly frustrated residues shown in red and minimally frustrated residues in green. In the 139-147 dimer, the proportion of frustrated residues (red line) increases with higher electrostatic force, resembling the trend observed in the 4BEX-assembly dimer.

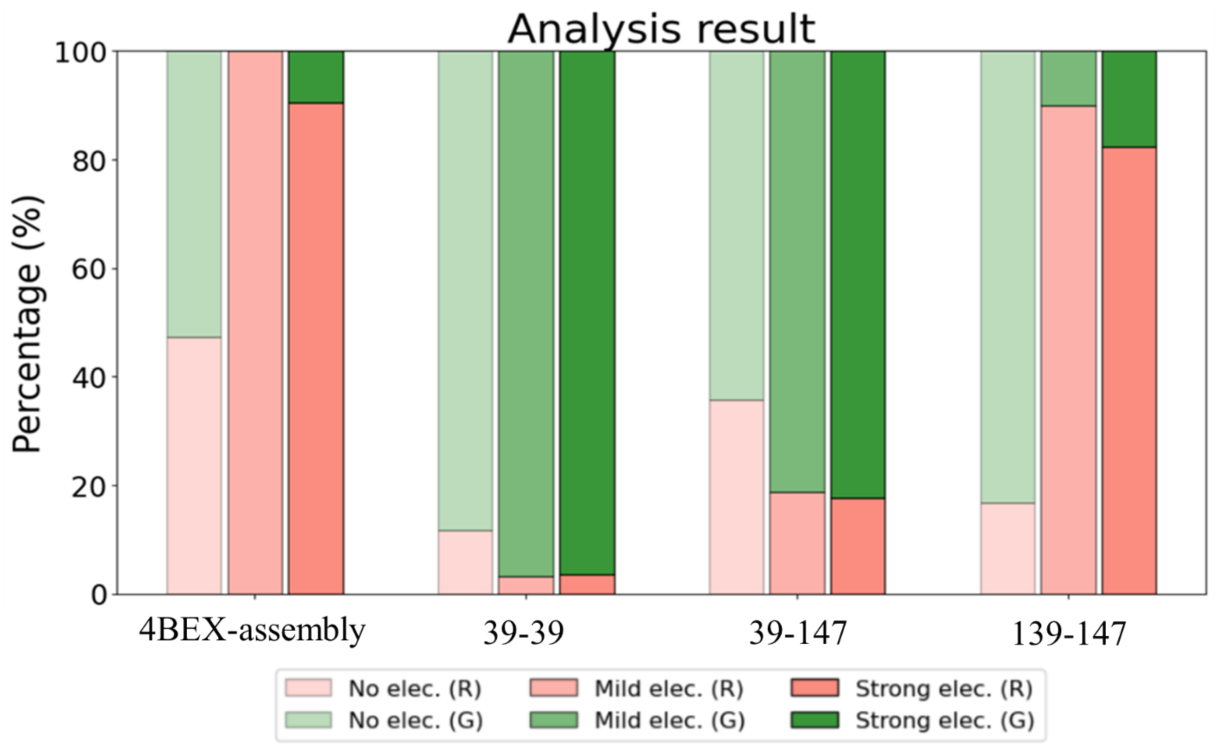


Figure S7 - Comparison of the four cofilin dimers' mutational frustration analysis is shown.

This histogram illustrates the percentage of frustrated interactions versus minimally frustrated interactions influenced by electrostatic forces for four cofilin dimers. Each column, from left to right, is labeled as "No electrostatic (No elec.)", "Mild electrostatic (Mild elec.)", and "Strong electrostatic (Strong elec.)", corresponding to the Debye-Hückel potential with dielectric constants $\varepsilon = \infty$, 80, and 33, respectively. Frustrated residues are depicted in red, while minimally frustrated residues are shown in green.
